# The nonlinear meccano of hyperactivity in Alzheimer

**DOI:** 10.1101/2023.10.09.561541

**Authors:** Giulio Bonifazi, Celia Luchena, Adhara Gaminde-Blasco, Carolina Ortiz-Sanz, Estibaliz Capetillo-Zarate, Carlos Matute, Elena Alberdi, Maurizio De Pittà

**Affiliations:** Basque Center for Applied Mathematics, Bilbao, Spain; Department of Neurosciences, University of the Basque Country, Leioa, Spain; Krembil Research Institute, University Health Network, Toronto, ON, Canada; Achucarro Basque Center for Neuroscience, Leioa, Spain; Centro de Investigación Biomédica en Red en Enfermedades Neurodegenerativas (CIBERNED), Leioa, Spain; Department of Physiology, University of Toronto, Toronto, ON, Canada

## Abstract

The pathophysiological process of Alzheimer’s disease (AD) is believed to begin many years before the formal diagnosis of AD dementia. This protracted preclinical phase offers a crucial window for potential therapeutic interventions, yet its comprehensive characterization remains elusive. Accumulating evidence suggests that amyloid-β (Aβ) may mediate neuronal hyperactivity in circuit dysfunction in the early stages of AD. At the same time, neural activity can also facilitate Aβ accumulation through intricate feed-forward interactions, complicating elucidating the conditions governing Aβ-dependent hyperactivity and its diagnostic utility. In this study, we use biophysical modeling to shed light on such conditions. Our analysis reveals that the inherently nonlinear nature of the underlying molecular interactions can give rise to various modes of hyperactivity emergence. This diversity in the mechanisms of hyperactivity may ultimately account for a spectrum of AD manifestations.

## Introduction

The clinical course of Alzheimer’s disease (AD) starts with the appearance of the first symptoms of mild cognitive impairment (MCI) (Morris, Storandt, et al., 2001). These symptoms then slowly progress to dementia as deficits emerge in multiple cognitive domains that are severe enough to produce loss of function (Jack Jr, Knopman, Jagust, Shaw, et al., 2010). Well-known neuropathological correlates of the disease are extracellular amyloid-β (Aβ) accumulations and intracellular depositions of neurofibrillary tangles in association with neurodegeneration by neuronal and synaptic loss reflected by progressive brain atrophy (Blennow and Zetterberg, 2018; Olsson et al., 2016; Huijbers et al., 2015; Jack Jr, Albert, et al., 2011).

The progression from MCI to dementia cannot be reverted at present, making AD intractable and the most common cause of dementia in elderly people (Kelley et al., 2015). On the other hand, Aβ and neurofibrillary tangles buildup could start long before MCI onset (Jack Jr, Knopman, Jagust, Shaw, et al., 2010; Jack Jr, Lowe, et al., 2009), pinpointing the existence of a preclinical phase of AD that most likely sets its fate, namely if, and to what extent, AD clinical features will develop (Sperling et al., 2011). Thus, to predict the risk of developing dementia by Alzheimer’s, characterizing the preclinical phase of the disease is crucial.

Current biomarker models of AD’s preclinical phases do not effectively predict the clinical syndrome of AD (Sperling et al., 2011). Aβ accumulation, for example, is recognized as a key early biomarker in AD etiology that is necessary yet likely not sufficient to incite the downstream pathological cascade of the disorder (Jack Jr, Knopman, Jagust, Shaw, et al., 2010; Frisoni, Altomare, et al., 2022). Hence, efforts are in the direction of identifying additional biomarkers that could predate Aβ accumulation or that, in combination with it, could predict the risk of developing the disease’s clinical syndrome reliably (Frisoni, Altomare, et al., 2022; Frisoni, Molinuevo, et al., 2020).

Several lines of evidence indicate that neuronal hyperactivity could also be a harbinger of AD-related dementia (Zott and Konnerth, 2022). Functional imaging studies in individuals with prodromal AD such as MCI reveal increased neuronal activity in the hippocampus and some neocortical areas (Huijbers et al., 2015; Mormino et al., 2012; Quiroz et al., 2010; Dickerson et al., 2005; Bookheimer et al., 2000), and those individuals often suffer from epileptic seizures possibly resulting from such excessive neuronal activation (Vossel, Beagle, et al., 2013; Palop and Mucke, 2009). Significantly, reducing hippocampal hyperactivation in MCI patients using the antiepileptic drug levetiracetam can partially restore cognitive function, especially in those patients also suffering from epileptic seizures (Vossel, Ranasinghe, et al., 2021; Bakker, Albert, et al., 2015; Bakker, Krauss, et al., 2012).

Experiments in AD-related mouse models suggest that high extracellular glutamate levels could cause prodromal neuronal hyperactivation. The presence of soluble Aβ in the extracellular space in the early stages of AD can reduce expression of astrocytic GLT1 transporters (Scimemi et al., 2013; Zott, Simon, et al., 2019; Hefendehl, LeDue, et al., 2016). Since these transporters are the main ones responsible for the clearance of extracellular glutamate, a reduction in their expression would result in a decreased uptake accounting for extracellular glutamate buildup (Rothstein et al., 1996).

Because Aβ production is activity-dependent (Cirrito et al., 2005; Kamenetz et al., 2003), so is Aβ-dependent GLT1 reduction, and the resulting glutamate buildup (Zott, Simon, et al., 2019). More extracellular glutamate then increases neuronal firing, which, in turn, promotes further glutamate release from synaptic terminals and Aβ production. In this fashion, a positive feedback loop is in place whereby initially low extracellular levels of Aβ and glutamate could be increased by ongoing neural activity and, in turn, increase the latter exacerbating AD clinical progression (Zott and Konnerth, 2022; Zott, Simon, et al., 2019).

The activity requirements of this feedback loop, however, are not understood. Preexisting baseline neural activity is necessary to promote Aβ-dependent hyperactivity (Zott, Simon, et al., 2019), but what levels of such baseline activity could hasten the latter are unclear (Zott and Konnerth, 2022). At the same time, the fact that not all AD patients with Aβ pathology develop seizures (Larner and Doran, 2006), nor all individuals with Aβ-correlated seizures develop AD (Mackenzie and Miller, 1994), suggests that the combination of Aβ buildup with neuronal hyperactivation predating the clinical phase of AD is variegated. Here, we explore this hypothesis, building a mathematical model of Aβ-dependent hyperactivity that considers the time-dependent expression of possible biomarkers associated with the phenomenon, such as extracellular Aβ and glutamate concentrations, and neuronal firing (Blennow and Zetterberg, 2018; Carter et al., 2019; Vossel, Tartaglia, et al., 2017; Busche and Konnerth, 2016). To model biomarkers’ temporal evolution, we consider three critical molecular pathways underpinning Aβ-dependent hyperactivity (Zott and Konnerth, 2022): (i) glutamate-mediated neuronal activity, (ii) Aβ-dependent glutamate uptake, and (iii) activity-dependent Aβ production (Figure 1a). The molecular reactions mediating such pathways and their interactions are nonlinear. Hence, a change in biomarker expression ensuing from a perturbation of one pathway is generally not proportional to that perturbation, nor can the singly perturbed pathway be accounted for. Instead, it is the result of combining the latter with the other interacting pathways. Moreover, the interactions among the different molecular pathways can mediate multiple positive feedback loops (Figure 1b). In such a scenario, the theory of non-linear dynamical systems predicts that multiple biomarker expressions could co-exist for the same preclinical stage of AD (Pisarchik and Feudel, 2014). We interpret this possibility by the existence of multiple trajectories toward clinical AD manifestations, each possibly associated with a specific risk of developing MCI and dementia.

**Figure 1.**
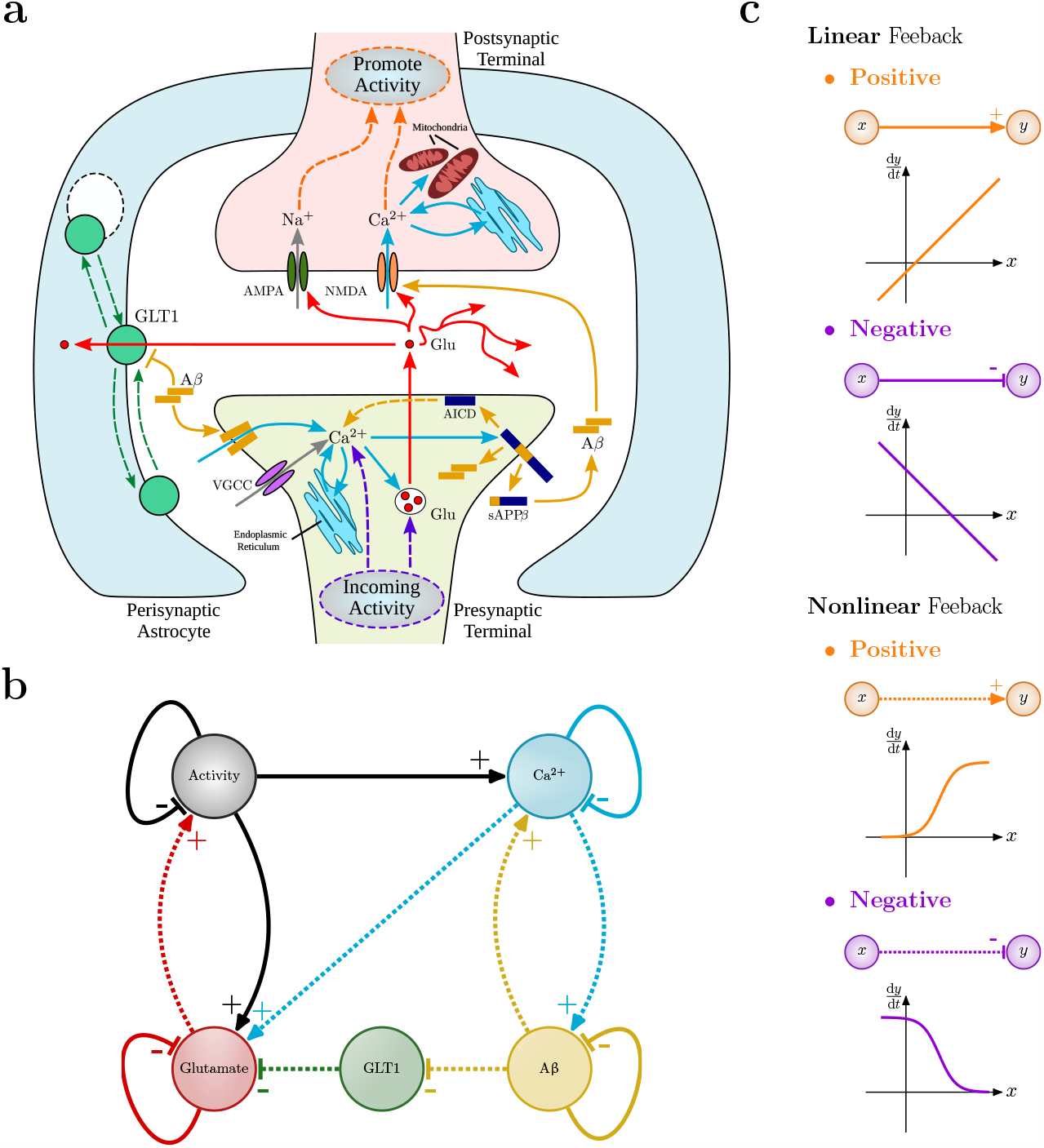
Aβ-dependent regulation of extracellular glutamate. **a** Molecular pathways regulating extracellular glutamate in the presence of Aβ accumulation. In healthy tissue, astrocytic glutamate transporters (GLT1) take up the majority of synaptically-released glutamate, shaping postsynaptic currents and regulating local neural activity (Tzingounis and Wadiche, 2007; Danbolt, 2001). GLT1 expression, however, can change with extracellular Aβ in a diverse fashion that depends on neural intracellular Ca^2+^ (Zott and Konnerth, 2022; Scimemi et al., 2013). Intracellular Ca^2+^ can elicit the secretory cleavage of the amyloid precursor protein (sAPP) both by activity-dependent and independent mechanisms (Bezprozvanny and Mattson, 2008). sAPP monomers are known to aggregate into soluble Aβ oligomers forming insoluble fibrils that finally deposit as amyloidogenic plaques (Viola and Klein, 2015). At the same time, Aβ and intracellular APP domains (AICD) can promote Ca^2+^ signaling through various intracellular and extracellular pathways (Gallego Villarejo et al., 2022). **b** Model of extracellular glutamate homeostasis including Aβ-dependent GLT1 regulation. Such regulation is by the interaction of activity-dependent glutamate release with Ca^2+^-dependent Aβ production, which is mediated by multiple loops of nonlinear interactions illustrated in **c**.

## Results

### Uptake by astrocytic transporters is the limiting process in extracellular glutamate clearance around Aβ accumulations

Aβ-dependent hyperactivity in preclinical AD is rarely whole-brain (Zott and Konnerth, 2022; Vossel, Beagle, et al., 2013). Instead, it is distributed and co-localizes microscopically with extracellular Aβ-depositions, also known as plaques, which are an early and predictive marker for the possible progression of preclinical to symptomatic AD (Jagust, 2018; Morris, Roe, et al., 2009). Before plaque deposition, however, soluble Aβ must accumulate at the plaque site, hastening plaque deposition (Hefendehl, Wegenast-Braun, et al., 2011; Hong et al., 2011). At the same time, as the plaque forms and grows, soluble Aβ continuously binds and unbinds from its surface, generating a toxic microenvironment around the plaque site where neuronal hyperactivity emerges (Busche, Chen, et al., 2012; Busche, Eichhoff, et al., 2008).

Imaging of synaptically evoked extracellular glutamate transients around Aβ plaques hints at reduced glutamate clearance rates by astrocytic transporters as a putative biophysical correlate for the toxic nature of the plaque microenvironment. It is helpful to understand how this happens in terms of the physical laws governing extracellular glutamate signaling. To this extent, we consider a tissue ball centered at an Aβ plaque as a model of the tissue microenvironment around the plaque (Figures 2a,b). How large our ball is in radius *R* will depend on the Aβ gradient under consideration, reflecting the extent of the plaque’s toxic microenvironment (Hefendehl, LeDue, et al., 2016). Typical radii are in the range of tens of micrometers (Hefendehl, LeDue, et al., 2016; Hefendehl, Wegenast-Braun, et al., 2011; Querol-Vilaseca et al., 2019; Pickett et al., 2016), thus much larger than individual synapses whose maximum dimension usually is of the order of tens of nanometers (Figures 2c,d) (Curran et al., 2021; Shapson-Coe et al., 2021; Kasthuri et al., 2015).

**Figure 2.**
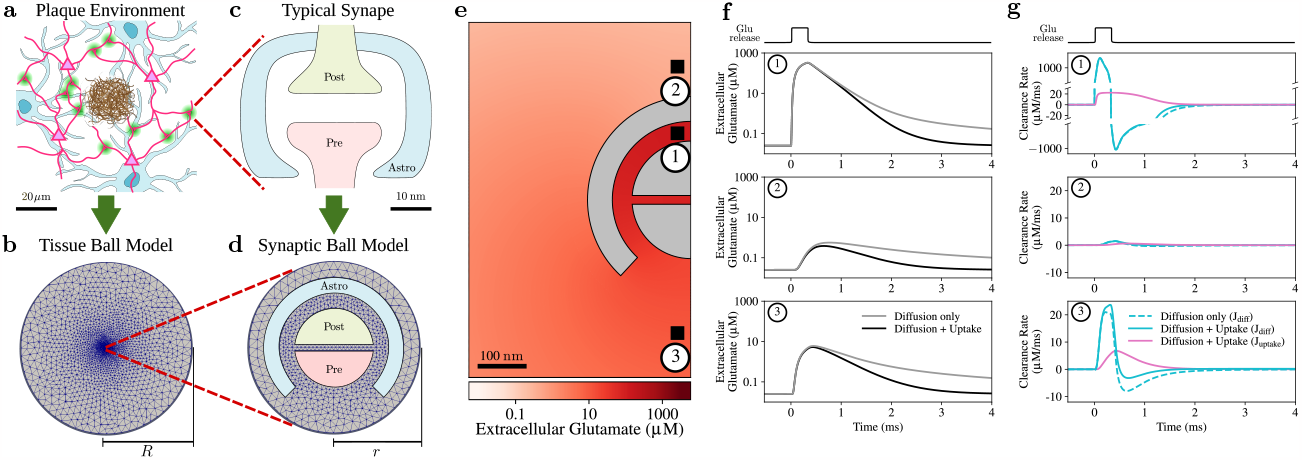
Contribution of astrocytic transporter uptake to extracellular glutamate clearance. **a,b** Model of the plaque’s microenvironment. The microenvironment where Aβ deposit is described by a tissue ball of radius *R* containing multiple synapses and astrocytes. **c,d** Typical synapses in this ball have much smaller radial size *r* and consist of pre- and postsynaptic terminals ensheathed by GLT1-expressing astrocytic processes. **e** Snapshot of the spatial distribution of perisynaptic glutamate taken 300 μs after a pulsed glutamate injection at the synapse center. **f** Associated glutamate time course at three sample points in the perisynaptic space (*black traces*). Compared with the time course in the absence of GLT1 uptake (*gray*), it may be appreciated how the uptake by astrocyte transporters reduces the transient duration of extracellular glutamate excess. **g** Considering the time evolution of the glutamate clearance by diffusion only vs. diffusion plus uptake reveals how diffusion is only marginally affected by uptake. That is, uptake by astrocytic transporters, rather than diffusion, modulates how fast extracellular glutamate is cleared.

Our tissue ball will generally comprise multiple cell bodies, dendrites, synapses, and astrocytic processes that are part of active neural circuits. We can characterize the geometry of the extracellular space associated with those circuits by the average fraction (*α*) of extracellular volume with respect to the ball volume. We also consider the average shape of the extracellular space in the ball as reflected by the tortuosity (*λ*) of the path of extracellular molecules diffusing around cellular obstructions created by the neuropil structure in the ball. The advantage of introducing these quantities is to be able to describe extracellular glutamate in time (*t*) and space (**x**) in the plaque microenvironment, i.e., *g* (**x**, *t*), by the macroscopic balance of synaptic release, *J*_syn*−*rel_(*α*), with clearance by passive diffusion, *J*_diff_ (*g, λ*), and active uptake by astrocytic transporters, *J*_uptake_(*g, α*) (Bergles, Diamond, et al., 1999). Namely (Syková and Nicholson, 2008)

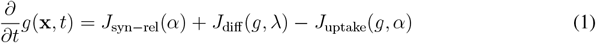

It is enlightening to look at solutions of the above equation in the surroundings of what a typical synapse could look like in our tissue ball (Lehre and Rusakov, 2002; Rusakov and Kullmann, 1998) (Figure 2e). Consider, for example, a brief glutamate injection at the center of the synaptic cleft mimicking synaptic release for 0 *< t ≤* 0.3 ms (near point “1” in Figure 2e). Equation 1 predicts a redistribution of this initial glutamate surge from the cleft to the extrasynaptic space by diffusion. The rise of extracellular concentration at any point in space is location-dependent due to synaptic and astrocytic obstacles (Figure 2f). On the other hand, for *t >* 0.3 ms, clearance of extracellular glutamate is qualitatively similar everywhere, either in the presence of or without uptake by astrocytic transporters (Figure 2f, dark vs. light gray traces).

The existence of uptake by transporters shortens the time course of extracellular glutamate. However, looking at individual mechanisms setting such time course (Figure 2g), it may be appreciated how glutamate uptake mainly contributes to the initial phase of glutamate clearance, and only when the local glutamate concentration is high enough. This follows from the sigmoid non-linearity 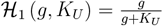 rising from the Michaelis-Menten-type kinetics of glutamate uptake (Tzingounis and Wadiche, 2007; Rusakov and Kullmann, 1998), whereby

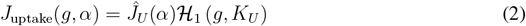

Equation 2 tells us that *J*_uptake_ is nonnegligible only when glutamate concentration approaches or exceeds the transporter’s affinity *K*_*U*_, that is, approximately, when 0.1*K*_*U*_ *< g <* 0.5*K*_*U*_ . When extracellular glutamate concentration largely exceeds *K*_*U*_ instead, e.g., *g >* 10*K*_*U*_, *J*_uptake_ saturates to the maximum uptake rate *Ĵ*_*U*_ (*α*). In this fashion, the maximal rate of glutamate clearance ensuing from diffusion and uptake cannot exceed *J*_diff_ (*g, λ*) + *Ĵ*_*U*_ (*α*). Therefore, since uptake but not diffusion changes with Aβ (Hefendehl, LeDue, et al., 2016), we conclude that uptake by astrocytic transporters is the mechanism that sets the limit for the shortest possible time course of extracellular glutamate in the synaptic cleft and extrasynaptically in the plaque microenvironment.

### The nonlinear nature of glutamate uptake results in nonuniform activity-dependent regulation of extracellular glutamate in the Aβ plaque microenvironment

The existence of a maximum clearance rate also implies that a maximum rate of glutamate supply to the extracellular space must exist beyond which glutamate starts accumulating extracellularly. This could happen, for example, when many synapses in our tissue ball are active for a protracted period and can be accounted for by a glutamate supply, *J*_syn*−*rel_ in equation 1, that is proportional to the neural activity rate (Supplementary Text Section **??**). Then, glutamate supply will exceed clearance for sufficiently large activity rates, promoting extracellular glutamate accumulation.

The activity rate promoting extracellular glutamate buildup will depend on the maximum uptake rate (*Ĵ*_*U*_). Astrocytic excitatory amino acid transporters – EAAT2 in humans and GLT1 in murine tissue – account for the majority of glutamate uptake in the adult brain, allowing neglecting the contribution to uptake by other transporter types (Tzingounis and Wadiche, 2007; Danbolt, 2001; Bergles and Jahr, 1998). We can thus assume that all transporters equally contribute to glutamate uptake in our tissue ball and estimate the maximum uptake rate by the product of the single transporter uptake rate (*r*_GLT1_) by the average transporter concentration (*n*(*α*)) found in our tissue ball. This concentration, and so the associated uptake rate, is the highest in the healthy tissue (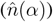) while progressively reducing with Aβ accumulation (Scimemi et al., 2013; Hefendehl, LeDue, et al., 2016). We can conveniently express this fact by the proportionality law

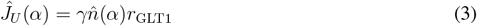

where the healthy tissue scenario corresponds to *γ* = 1 when the maximum uptake rate is the fastest because transporter expression is the largest, namely, 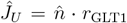 . Conversely, when 0 *≤ γ <* 1, a decrease in transporter expression and thus in uptake is envisaged by the presence of extracellular Aβ and we look at this scenario for the emergence of toxic extracellular glutamate accumulations.

In Figures 3a,b, we consider extrasynaptic glutamate dynamics for different transporter expression at two sample rates (*ν*) of synaptic release in the gamma frequency range that could be representative of cognitive-relevant neural activity in our tissue ball (Fries et al., 2007; Buzsaki, 2006). On the upper end of the spectrum of transporter expression, when *γ* = 1, our simulations predict that extrasynaptic glutamate will generally transiently accumulate in the range of 1 150 μM (pink traces in Figure 3b). Though considerably higher than resting extracellular glutamate levels reported to be *<* 0.1 μM (Herman and Jahr, 2007; Cavelier and Attwell, 2005), these concentrations agree with estimates of non-toxic physiological glutamate levels surrounding active synapses (Lehre and Rusakov, 2002; Barbour, 2001; Clements et al., 1992). Conversely, on the lower end of the uptake spectrum, in conditions of strongly reduced transporter expression (*γ* = 0.1), we find instead dangerously high glutamate concentrations, i.e., *>* 200 μM (dark red traces in Figure 3b). Glutamate concentrations of such magnitude are estimated to mediate acute and chronic excitotoxicity (Lewerenz and Maher, 2015).

**Figure 3.**
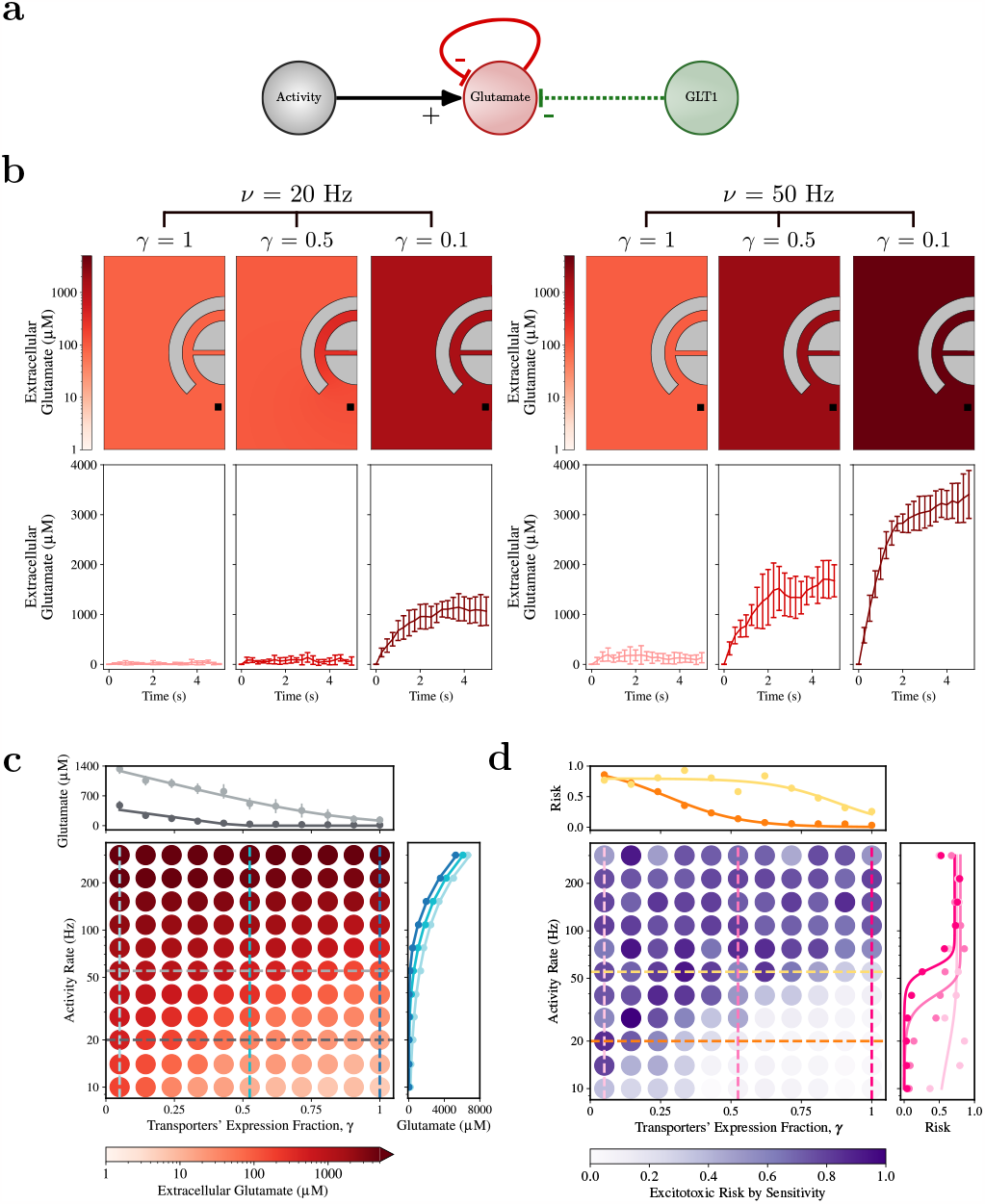
Nonlinear glutamate accumulation. **a** Model setup to characterize how activitydependent synaptic release influences extracellular glutamate clearance. **b** Simulated perisynaptic glutamate levels for Poisson-distributed glutamate pulse injections at the synapse center at two rates (*ν*) representative of sensory-relevant synaptic activity and for different expressions of astrocytic glutamate transporters (*γ*). The *top* panels are snapshots of extracellular glutamate at *t* = 5 s around a typical microenvironment synapse, averaged over *n* = 20 simulations. The *bottom* panels show the average glutamate time course (± s.t.d.) at the location marked by the *black square* in the top panels. **c** Average steady-state extracellular glutamate in the plaque microenvironment as a function of *γ* and *ν*. (*Insets*) Representative curves for steady-state glutamate concentrations attained by fixing *γ* or *ν* show a nonlinear behavior marked by an inflection point beyond which glutamate can increase toward potentially toxic levels. **d** Computation of the derivative of glutamate concentration with respect to *ν* at fixed *γ* provides a measure of how susceptible glutamate is to increase in response to rate variations and thus reflects the risk of developing such toxic glutamate accumulations. (*Insets*) Tracing this risk as a function of *ν* or *γ* reveals a sigmoid curve, whose inflection point nonuniformly changes with *γ* (respectively, *ν*).

We observe healthy and toxic microenvironments for the same transporter expression (*γ* = 1 and *γ* = 0.1, respectively) regardless of the activity rate under consideration. However, this would not be the case for an intermediate (*γ* = 0.5) reduction of transporter expression. In such a scenario, it may be appreciated how glutamate reaches physiological concentrations *≤* 150 μM for *ν* = 20 Hz but builds up to toxic levels *>*1500 μM that are 10-fold larger, for just a 2.5-fold increase in activity when *ν* = 50 Hz. This asymmetric increase of extracellular glutamate with respect to the activity rate follows from the nonlinearity of equation 1 introduced by the transporter kinetics, whereby the glutamate uptake rate saturates for 20 *< ν <* 50 Hz. Thus, at *ν* = 50 Hz, but not at 20 Hz, glutamate supply (*J*_syn*−*rel_) exceeds clearance, allowing for the quick buildup of toxic concentrations.

The asymmetric nature of activity-dependent glutamate accumulation not only tells us that the activity requirements to develop toxic glutamate accumulations in the plaque microenvironment change with different transporter expressions but also that the way they change and the associated risk of glutamate toxicity, are nonuniform. We can appreciate such a nonuniformity, mapping the average extracellular glutamate concentration in our tissue ball (Figure 3c) and its derivative (Figure 3d) as functions of the activity rate for multiple transporter expressions. It may be seen how toxic glutamate concentrations can only be attained beyond a threshold activity rate that varies with transporter expression (Figure 3c, right inset). Such a threshold can be inferred from the sigmoid curve fitting the estimated derivative (Figure 3d, right inset). Additionally, the fact that such derivative is sigmoid hints that extracellular glutamate nonlinearly increases with increasing baseline synaptic activity, as would be the case for emerging Aβ-dependent hyperactivity (Zott, Simon, et al., 2019). The glutamate increase indeed approaches zero for small increments of synaptic release at low baseline activity but progressively grows towards a maximum when such baseline increases. We can also regard the rate of change in glutamate buildup for increasing synaptic activity as an estimation of the sensitivity, and thus the risk (Frey and Sumeet, 2002), for those buildups to grow toxic. In this framework, the risk of developing glutamate toxicity will nonuniformly change along the sigmoid derivative by the dynamic modulation of the release threshold as transporter expression reduces with accumulating Aβ (Scimemi et al., 2013; Tong et al., 2017).

### The threshold for glutamate-mediated neural hyperactivation is complex

In parallel with transporter expression’s Aβ-dependent dynamics, an additional dynamical component in the modulation of the activity threshold for the onset of glutamate toxicity comes from the possible time-dependence of synaptic glutamate release by spontaneous or evoked fluctuations in the neural activity in the plaque microenvironment. We can think of this activity as the result of afferent excitation from the outside of the microenvironment (*ν*_ex_) in combination with local firing activity (*ν*) resulting from the balance of two opposite feedback mechanisms (Figure 4a). One is the positive feedback of extracellular glutamate on neural activity and the synaptic glutamate release associated with it (Zott, Simon, et al., 2019), which we denote by some generic function *F*_syn*−*rel_(*g*) for the moment. The other is the negative feedback, likely mediated by multiple molecular pathways that homeostatically maintain a baseline activity (*ν*_0_) in our tissue ball (Frere and Slutsky, 2018). The interplay of these mechanisms’ results can be described by the differential equation

**Figure 4.**
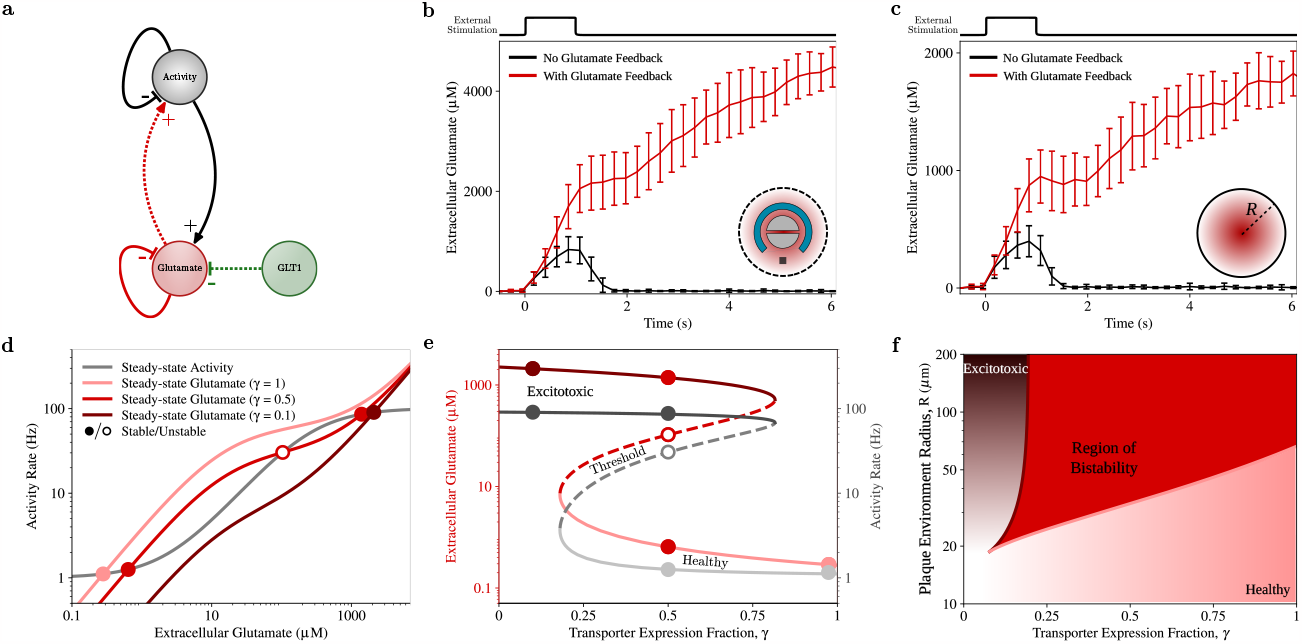
Diverse conditions for excitotoxicity. **a** Model of glutamate clearance including the positive feedback of glutamate on neural activity. **b** Simulated glutamate time course in the perisynaptic space and **c** in the whole plaque microenvironment for 1 s-long activity pulse. The positive feedback of glutamate on neural activity promotes synaptic glutamate release, which can quickly exacerbate toxic glutamate build-ups. **d** Graphical analysis of the steady-state glutamate concentrations and activity rates in the plaque microenvironment reveal how healthy and excitotoxic conditions could be both plausible for intermediate astrocytic transporter expressions *γ* (solid red circles) in the presence of glutamate feedback on the activity. **e** Compatibly with this scenario, the bifurcation diagrams for the steady-state glutamate/activity as a function of *γ* reveal the existence of a region of bistable glutamate and activity levels for 0.2 *< γ <* 0.8. **f** In such a region, the threshold of excitotoxicity nonlinearly varies with *γ* and radial size *R* of the plaque microenvironment.

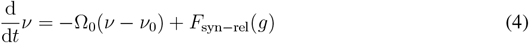

The first right-hand side term in the above equation reflects the homeostatic feedback, which we can consider, without loss of generality, to be linear with the difference of the instantaneous local activity rate from baseline, i.e., *ν − ν*_0_, by the homeostatic recovery rate constant Ω_0_ (O’Leary and Wyllie, 2011). However, that is not true for the positive feedback term *F*_syn*−*rel_. The fact that neurons can only fire when depolarized beyond a firing threshold by excitatory (glutamatergic) synapses (Rauch et al., 2003) implies that the feedback of extracellular glutamate on neuronal depolarization, and thus firing, and downstream synaptic release, kicks in only around and beyond a threshold glutamate concentration. The existence of such a threshold may conveniently be described by a sigmoid glutamate-dependent change of the activity rate such as (Supplementary Text Section **??**)

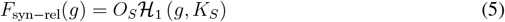

where *K*_*S*_ is the threshold glutamate concentration for feedback emergence, and *O*_*S*_ is the maximum change rate as extracellular glutamate grows large beyond the threshold, e.g., *g >* 10*K*_*S*_.

We emphasize the sigmoid nonlinearity introduced by the glutamate-dependent feedback because, together with the nonlinearity of transporters’ uptake (equation 2), it could put forth a molecular switch for the onset of glutamate toxicity. We may appreciate this switch in Figures 4b,c which show representative numerical solutions of equation 1 for the time course of the average extracellular glutamate following a transient increase of afferent activity (*ν*_ex_, top pulses, for 0 *≤ t <* 1 s), respectively around a representative synapse of our tissue ball, and in the whole ball. In the absence of feedback, i.e., *F*_syn*−*rel_ = 0, glutamate builds up during the transient activity increase but is then quickly removed by diffusion and uptake, decreasing to pre-stimulus concentrations (*black traces*). In the presence of feedback instead, glutamate grows higher during and after stimulation, eventually switching to persistent toxic concentrations (*red traces*). That is, the positive feedback of glutamate on neural activity amplifies the initial glutamate increase by afferent stimulation, which drives a vicious cycle of hyperactivation associated with toxic glutamate accumulation (Zott and Konnerth, 2022; Lewerenz and Maher, 2015). The feedback’s sigmoid nature is such that as glutamate accumulates promoting activity rates, the associated increase in synaptic release quickly grows beyond the maximum clearance capacity by diffusion and uptake, resulting in persistent glutamate accumulation.

The persistent glutamate accumulation has a simple mathematical interpretation that can be evinced from the graphical solutions of equations 1 and 4 for the steady state. Figure 4d shows these solutions by the intersection of the gray and red curves respectively for 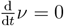 and 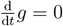 for the three transporter expressions (*γ*) already considered in Figure 3. One solution exists at low glutamate and activity rates for large transporter expression (*γ* = 1, pink circle), reflecting a healthy tissue state. Likewise, one solution exists in the toxic tissue state characterized by high glutamate and activity occurring for strongly reduced transporter expressions (*γ* = 0.1, dark red circle). However, both healthy and toxic states are viable for intermediate transporter expressions (*γ* = 0.5, full red circles) consistent with a scenario of bistability. In this scenario, as revealed by our simulations in Figure 4c, we could start from the healthy state but end in toxicity as soon as activity and glutamate increase beyond levels set by the unstable solution marked by the empty red circle.

Mapping unstable and stable solutions for all possible transporter expressions results in the diagrams in Figure 4e, technically known as bifurcation diagrams. Their characteristic ‘S’ shape is a hallmark of bistability (Slepchenko and Terasaki, 2004) insofar as healthy and toxic states, respectively represented by the diagrams’ low (pink) and high branches (red), coexist for transporter expressions between 0.2 *< γ <* 0.8. For such expression, the plaque microenvironment could thus become toxic or stay healthy depending on whether the local activity and glutamate levels are above or below the dashed threshold. At the same time, as far as the plaque microenvironment exists in the bistable regime, switching between healthy and toxic conditions is always possible by appropriate perturbations of activity and glutamate across the threshold. This can be therapeutically advantageous since toxicity could be reverted, thus slowing, halting, or reverting AD progression.

Because the bistability region separates between healthy-only (*γ >* 0.8) and toxic-only transporter expressions (*γ <* 0.2), we can liken it to a threshold for the transition between the two as transporter expression reduces with Aβ accumulation during preclinical AD progression. However, the nature of such a threshold is complex insofar as the emergence of bistability does not necessarily cause the transition but rather introduces the chance for it. Since this chance correlates with how far the dashed threshold in the bifurcation diagrams is with respect to the final state (Izhikevich, 2007), we conclude that the *risk* of toxicity turns into certainty when transporters reduce below *γ ≈* 0.2. Conversely, the *opportunity* to rescue healthy conditions increases with transporter expressions approaching the upper boundary of the bistability region, i.e., *γ ≈* 0.8.

We want to understand when it makes sense to look for bistability for diagnostics and therapeutic purposes. Since the spatial extent of Aβ accumulations controls how far toxicity develops from them (Hefendehl, LeDue, et al., 2016), toxic conditions could then be envisaged only for sufficiently large tissue balls. Indeed, mapping the boundaries of healthy (pink) and toxic conditions (red) in terms of the microenvironment’s transporter expression and radius (*R*) in Figure 4f reveals how they delimit a hashed region of bistability originating from a cusp. Below this cusp, we are looking at small tissue balls of radii *R <* 20 μm where locally-released glutamate can always diffuse out from regardless of their small transporter expression (*γ <* 0.1) (white shades). Hence, there is no clear separation between healthy and toxic states in such confined tissue environments because toxic glutamate concentrations can only build up by appropriate levels of exogenously maintained local activity (Supplementary Figure **??**). Conversely, as we look at larger tissue balls (*R >* 20 μm), locally released glutamate molecules need to travel longer distances to escape from it, facilitating glutamate accumulation regardless of transporter expression. This increases the chance of developing toxicity as reflected by a bistability’s transporter expression range that increases with the ball’s radius.

It is intriguing to correlate the growth of the bistability region with its robustness against local variations in transporter expressions (Scimemi et al., 2013; Hefendehl, LeDue, et al., 2016; Tong et al., 2017) as the spatial extension of Aβ deposition increases with AD progression. Insofar as the gross of this deposition likely predates the onset of clinical AD (Jack Jr, Knopman, Jagust, Shaw, et al., 2010), we could predict that the closer to clinical manifestations we are, the more we could exploit bistability to avoid toxic developments that would exacerbate AD symptoms. In practice, however, we will have to take into account that the degree of Aβ deposition also correlates with alterations in the tissue cytoarchitecture accounted for by variations of the extracellular volume fraction (*α*) and the tortuosity of the diffusion pathway of glutamate (*λ*, equation 1) (Bondareff, 2013; Syková, Voříšek, et al., 2005). Although such variations may be variegated, they will ultimately act against the emergence of bistability for sufficiently high Aβ depositions, regardless of their underpinning molecular mechanisms (Supplementary Figure **??**).

### Multiple trajectories with unique risk exist towards clinical AD syndrome

The process of Aβ accumulation preluding clinical AD is inherently dependent on intracellular Ca^2+^ and vice versa (Bezprozvanny and Mattson, 2008). Altered neuronal cytosolic Ca^2+^ accelerates Aβ formation, whereas Aβ peptides, particularly in soluble oligomeric forms, induce Ca^2+^ disruptions. This reciprocal interaction results in a feed-forward cycle of toxic Aβ generation and Ca^2+^ perturbations, which can exacerbate AD pathology (Demuro et al., 2010). Moreover, another feed-forward cycle exists between Ca^2+^ and neural activity. Ongoing activity modulates intraneuronal Ca^2+^ levels (Higley and Sabatini, 2012), while in turn Ca^2+^ crucially regulates glutamate release from synaptic terminals (Schneggenburger and Neher, 2005). In this way, the Ca^2+^-dependent Aβ accumulation becomes inherently activity-dependent, and so does astrocytic transporter expression. How could the combination of these pathways influence the onset of hyperactivity in prodromal AD?

The sigmoid laws governing the bistability of the glutamate vicious cycle on activity as a function of transporter expression (Figure 4) also account for bistability as a function of the tissue baseline activity (*ν*_0_) (Figure 5a,b) which is known to increase with AD emergence (Zott and Konnerth, 2022). In particular, our tissue model predicts the existence of a threshold of glutamatergic activity (dashed lines in Figure 5b.2) for basal activity rates up to about 15 Hz. Below this rate, it will always be possible to rescue the healthy state starting from excitotoxic conditions by perturbations of initial conditions below that threshold. Conversely, as soon as basal activity increases beyond 15 Hz, excitotoxicity becomes practically unavoidable (Figure 5b.3). A similar scenario of bistability also occurs for Ca^2+^-dependent Aβ accumulation (De Caluwé and Dupont, 2013, and Supplementary Figure **??**). The process can indeed be described by sigmoid laws akin to those governing the glutamate/activity vicious cycle (Supplementary Text Section **??**), predicting that low vs. high Aβ and Ca^2+^ concentrations coexist for the same basal activity rate up to about 50 Hz (Figure 5d).

**Figure 5.**
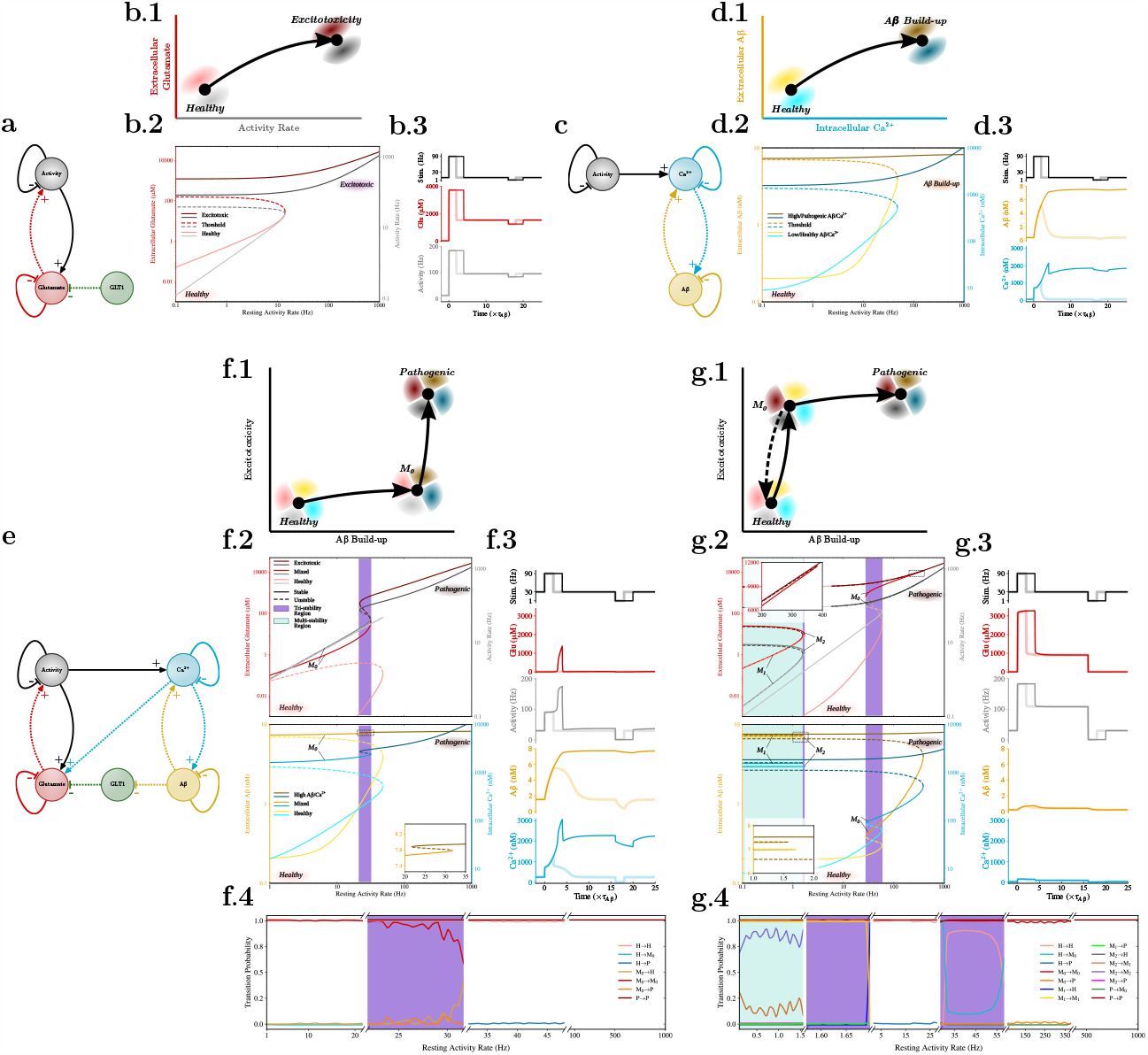
Different trajectories for AD emergence. Two activity-dependent feed-forward processes exist in Aβ pathogenesis respectively by **a,b** the glutamate-mediated vicious cycle on activity, and **c,d** the Ca^2+^-dependent vicious cycle on Aβ build-up. **b.2,c.2** Both cycles can mediate a scenario of bistability for basal activity rates with **b.3,c.3** irreversible pathogenic states. **e** The two cycles are also coupled by nonlinear Ca^2+^-dependent glutamate release and nonlinear Aβreduction of transporter uptake, accounting for multiple scenarios of AD emergence, depending on the biophysical properties of affected neural circuits. **f** In one scenario, Aβ predates excitotoxicity, whereas in another **g** excitotoxicity may anticipate Aβ build-ups. These different scenarios are possible by intermediate states manifesting only some pathological features (e.g., excitotoxic conditions vs. Aβ/Ca^2+^ build-ups), which can account for multiple nuances of pathogenic tissue in AD progression beyond simple distinction between healthy vs. pathological. **f.4,g.4** Probabiities of permissible transitions between states in the function of the basal activity rate (*ν*_0_). The probability of transition between two states *A* and *B*, i.e., *A → B*, is computed following (Kaszás et al., 2019) by the area ratio between the set of all initial conditions that from *A* ends in *B* for an increment *ν*_0_ *→ ν*_0_ +Δ, divided by the set of initial conditions ending in *B* before such increment.

The combination of the bistability scenarios of the two processes would account, at least in principle, for four possible tissue states: a healthy state characterized by low extracellular glutamate and local activity with low Aβ and Ca^2+^ levels, a pathogenic state where excitotoxicity coexists with high Aβ and Ca^2+^ concentrations, and two other “mixed” states where low glutamate/activity (respectively low Aβ/Ca^2+^) combine with high Aβ and Ca^2+^ (respectively, high glutamate and activity rates). However, because sigmoid laws govern how Ca^2+^ regulate synaptic glutamate release (Schneggenburger and Neher, 2005), and how Aβ could reduce glutamate uptake by astrocytic transporters (Figure 5e) (Fernández-Tomé et al., 2004, and Supplementary Text Section **??**) we should expect a nonuniform distribution for the probability of occurrence of one state vs. another. We thus set to characterize the admissible states in AD emergence by including in our model of glutamate-mediated hyperactivity an effective description of Ca^2+^dependent Aβ build-up (De Caluwé and Dupont, 2013, and Supplementary Text Section **??**) and considering two different scenarios of Ca^2+^ affinity of synaptic release (*K*_*R*_) and activity-dependent Ca^2+^ signaling (*η*_*C*_) that could be representative of the heterogeneity of cortical synaptic circuits (van Oostrum et al., 2023; Wang and Dudko, 2021; Zhu et al., 2018, and Supplementary Text Section **??**).

Figure 5f considers the case of synaptic circuits characterized by a moderate Ca^2+^ affinity (*K*_*R*_ = 3.6 μM) and an equally moderate activity-dependent Ca^2+^ production (*η*_*C*_ = 1 s). Our model predicts that healthy vs. pathogenic states in such circuits are separated by an intermediate one (M_0_) characterized by extracellular glutamate levels and activity rates close to healthy conditions while Aβ and Ca^2+^ concentrations approach pathogenic ones (Figure 5f.2). In this case, an increase in the local basal activity that results in excitotoxic pathogenic conditions could only be partially mitigated by any treatment to reduce basal activity. Reducing basal activity in such pathogenic conditions only rescues glutamatergic activity levels attained in healthy conditions, but not Aβ/Ca^2+^ ones (solid dark traces in Figure 5f.3). Thus, in agreement with the Aβ cascade hypothesis of AD (Selkoe and Hardy, 2016), this scenario predicts that Aβ build-up predates the onset of excitotoxicity.

The opposite scenario is instead observed in Figure 5g, which considers the case of a sample tissue whose local circuits could display increased synaptic release (*K*_*R*_ = 80 nM) but weaker activity-dependent Ca^2+^ production (*η*_*C*_ = 0.1 s). The bifurcation diagrams associated with this case reveal instead the existence of an intermediate state M_0_ where high glutamate/activity rates coexist with low Aβ/Ca^2+^ levels (Figure 5g.2). This suggests that such circuits appear more likely to develop excitotoxic conditions, as reflected by the robust glutamate and activity surge against a mild increase in Aβ/Ca^2+^ for the increase in basal activity that in the previous scenario was, on the contrary, primarily amyloidogenic. Moreover, it may be noted how reverting such an increase could revert both excitotoxic conditions and Aβ and Ca^2+^ build-ups, ultimately rescuing healthy conditions (Figure 5g.3).

A closer look at the bifurcation diagrams associated with the two tissue scenarios pinpoints the origin of such reversibility to the domain of existence of the diagram branch associated with the intermediate M_0_ state and how it is positioned between the healthy and pathogenic ones. In Figure 5f.2, the M_0_ branch exists for all basal activity rates up to the right boundary of the purple area, which occurs before the healthy branch’s termination. Moreover, the M_0_ and pathogenic branches together span the whole range of permissible basal activity rates, making rescuing healthy conditions impossible from either state by the sole reduction of the basal activity. Conversely, in Figure 5g.2, the M_0_ branch only exists in a confined range of basal activity values, originating around 25 Hz and extending beyond the healthy branch’s upper limit, up to roughly 450 Hz. In this way, any basal activity reduction below the purple-shaded area’s lower boundary could rescue healthy conditions.

The existence of “mixed” tissue states accounting for excitotoxicity but not Aβ/Ca^2+^ build up, or vice versa, is robust for a broad range of biophysical tissue parameters (Supplementary Figure **??**). At the same time, different biophysical properties could account for even more complex scenarios where more than one intermediate state exists between healthy and pathogenic (Supplementary Figure **??**). For example, the bifurcation diagrams in Figure 5g.2 also reveal that two additional states, M_1_ and M_2_, account for similar Aβ/Ca^2+^ build-ups but different intermediate levels of extracellular glutamate accumulation and associated activity at low basal activity rates (*<*1.5 Hz). An analysis of the permissible transitions allowed from/to these states (5g.4) then hints that transient increases of basal activity could be sufficient to destabilize such states (e.g., M_1_, M_2_*→* H), making the tissue fall back to somehow healthier conditions of lower glutamatergic activity and Aβ burden.

The chance to end in one of such intermediate states during AD emergence will ultimately depend on the initial state of the tissue. This is because the set of initial values of extracellular glutamate and Aβ concentrations, intracellular Ca^2+^, and local activity rates that determine whether the tissue ends up in one state or another change with the basal rate of activity (Figures 5f.4 and 5g.4). Thus, as the basal rate changes with the AD emergence, so does the probability of transitioning from healthy to intermediate states before ending into pathological ones. Since the tissue’s biophysical properties will ultimately dictate how this transition probability changes, we conclude that the risk of developing excitotoxicity vs. Aβ/Ca^2+^ build-ups in AD is variegated throughout the brain, with different sites expected to develop only some aspects of the pathology rather than others.

## Discussion

We presented a mathematical model of Aβ-dependent neuronal hyperactivation as an early-stage hallmark of neuronal dysfunction in Alzheimer’s disease (AD) (Zott, Simon, et al., 2019; Busche, Chen, et al., 2012; Busche, Eichhoff, et al., 2008). The model replicates the experimental observation that extracellular Aβ leads to the impairment of glutamate uptake by astrocytic transporters, resulting in the accumulation of perisynaptic glutamate. In turn, the excessive extracellular glutamate increases the activity levels further, mimicking neuronal depolarization by glutamate binding to ionotropic glutamate receptors, potentially creating a vicious cycle that initiates and sustains hyperactivity (Zott and Konnerth, 2022; Busche and Konnerth, 2016). In analogy with experimental observations, this vicious cycle depends on the baseline neuronal activity (Zott, Simon, et al., 2019). Neurons affected by the Aβ-dependent reduction of glutamate uptake can experience a buildup of glutamate levels, perpetuating hyperactivity. In contrast, inactive neurons with low levels of glutamatergic stimulation are less likely to become part of this harmful cycle.

The interaction of glutamatergic activity with intracellular Ca^2+^ and Aβ production makes the onset Aβ-dependent hyperactivation conditions inherently variegated. Neural hyperactivity generally positively correlates with Aβ levels and vice versa, mirroring experimental findings (Cirrito et al., 2005; Kamenetz et al., 2003; Bero et al., 2011). However, we also reveal that potentially excitotoxic hyperactivity and pathogenic Aβ buildups could be attained independently for distinct baseline neuronal activities, depending on the tissue’s biophysical properties. In other words, inherent differences in brain regions’ cytoarchitecture and molecular organization could account for different mechanisms of AD emergence, eventually reflecting on the heterogeneity of the disease at later, more advanced syndromic stages (Ten Kate et al., 2018; Zhang et al., 2016).

AD’s regional specificity almost certainly correlates with dynamical heterogeneity (Habes et al., 2020). Effectively, our model predicts that different tissue biophysical properties could result in different trajectories for AD emergence, where either hyperactivity predates Aβ deposition or vice versa. Moreover, both scenarios could be subjected to hysteresis, whereby the effect of a perturbation of baseline activity would generally vary with the AD progression stage. This supports the notion that AD etiology is complex, and the disease encompasses a spectrum of subtypes that could conveniently be stratified by region and stage (Young et al., 2018).

Our prediction that AD progression could be characterized by various mixed states where excitotoxicity emerges before Aβ accumulation, and vice versa, corroborates the idea of a continuum of AD pathology (Jack Jr, Knopman, Jagust, Shaw, et al., 2010; Jack Jr, Knopman, Jagust, Petersen, et al., 2013). At the same time, it also challenges our notion of a single preclinical stage of the disease in favor of multiple scenarios of AD emergence, each possibly characterized by a unique risk (Frisoni, Altomare, et al., 2022). In this framework, the fact that hyperactivity stemming from astrocyte glutamatergic dysfunction could predate Aβ accumulation is a promising avenue for biomarker development in the asymptomatic phase of the disease (Sperling et al., 2011; Carter et al., 2019). On the other hand, we also argue that, in light of the complex nonlinear nature of the interactions between excitotoxicity and Aβ/Ca^2+^ accumulations, the synergy between Aβ deposition and hyperactivity, rather than their additive effects, is likely a better predictor of AD emergence, as it is the case, for example, of considering Aβ plaques and tau tangles together in diagnostics of later stages of the disease (Pascoal et al., 2017).

The nonlinearity of the molecular biology underpinning AD pathology remains to be fully characterized. Future extensions of our model should also take into account the possibility of direct coupling of activity with Aβ production through endosomal and ectoenzymatic pathways, the scenario of direct modulation of GLT1 expression by Ca^2+^-dependent pathways in addition to Aβ (Todd and Hardingham, 2020; Stargardt et al., 2015), and the possibility that Aβ-mediated oxidative stress and gliosis could modify GLT1 expression by alterations of astrocytic cytoskeleton (Carter et al., 2019; Wyssenbach et al., 2016; Alberdi et al., 2013). The nonlinearity of these pathways and many possible others behind AD molecular pathophysiology (Henstridge et al., 2019), will only enrich the mosaics of potential tissue states in AD emergence (Latulippe et al., 2018), and, thus, the putative trajectories for the disease’s onset. Since the combination of the different nonlinearities uniquely characterizes the risk of each trajectory, interpreted as the chance for their occurrence, it also opens unforeseen avenues for personalized treatment by risk reduction interventions informed by the inherent nonlinear nature of the underpinning reactome (Frisoni, Altomare, et al., 2022; Frisoni, Molinuevo, et al., 2020).

In the search for effective AD theranostics, the question that stands out is which features or circumstances dictate that some neuronal cells and brain regions succumb to catastrophic fates when burdened by excitotoxic Aβ lesions, while other cells and regions seem to have a higher threshold of withstanding toxicity and manage to retain their normal function (Mrdjen et al., 2019). The modeling framework introduced in this study helps address this question, providing a mathematical characterization of the rules governing the complex interplay between intrinsic and extrinsic properties of this highly specific regional and cellular vulnerability to AD. Including this framework in current data-driven approaches for AD monitoring (Chen et al., 2021; Wang, Li, et al., 2021; Huang et al., 2020) will ultimately serve to refine diagnosis, help explain the molecular mechanisms of AD development and progression, and reveal potential compensatory mechanisms to bolster resilience.

## Methods

The detailed derivation of the models presented in this study, the exposition of the numerical methods adopted for their simulation and analysis, and the estimation of the models’ biophysical parameters may be found in the Supplementary Online Material.

## Acknowledgements

GB and MDP acknowledge the generous support of a Junior Leader Fellowship to MDP by ‘la Caixa’ Foundation (Grant ID: LCF-BQ-LI18-11630006). De Pittà’s lab is supported by a Seed Fund by the Krembil Foundation. CL, AGB, COS, ECZ, and EA are supported by MICIN/AEI (Grant ID: CPP2021-008389) and BIO22/ALZ/014 (Grant ID: PID2022-140236OB-I00) funded by BIOEF/The Basque Government. CM is supported by the Basque Government (Grant ID: IT-1551-22) and the CIBERNED-Instituto Carlos III (Grant ID: CB06/05/0076).

## Notes

### Competing Interest Statement

The authors have declared no competing interest.

